# Twenty five new viruses associated with the Drosophilidae (Diptera)

**DOI:** 10.1101/041665

**Authors:** Claire L. Webster, Ben Longdon, Samuel H. Lewis, Darren J. Obbard

## Abstract

*Drosophila melanogaster* is an important laboratory model for studies of antiviral immunity in invertebrates, and *Drosophila* species provide a valuable system to study virus host range and host switching. Here we use metagenomic RNA sequencing of *ca*. 1600 adult flies to discover 25 new RNA viruses associated with six different drosophilid hosts in the wild. We also provide a comprehensive listing of viruses previously reported from the Drosophilidae. The new viruses include Iflaviruses, Rhabdoviruses, Nodaviruses, and Reoviruses, and members of unclassified lineages distantly related to Negeviruses, Sobemoviruses and Poleroviruses, Flaviviridae, and Tombusviridae. Among these are close relatives of *Drosophila X virus* and *Flock House virus*, which we find in association with wild *Drosophila immigrans*. These two viruses are widely used in experimental studies but have not previously been reported to naturally infect *Drosophila*. Although we detect no new DNA viruses, in *D. immigrans* and *D. obscura* we identify sequences very closely related to *Armadillidium vulgare* Iridescent virus (Invertebrate Iridescent virus 31), bringing the total number of DNA viruses found in the Drosophilidae to three.

## Introduction

*Drosophila melanogaster* is an important model system for the study of antiviral immunity in invertebrates^1–4^, and has been instrumental in defining all of the major insect antiviral immune mechanisms, including the RNAi, IMD, Toll, autophagy, and Jak-Stat pathways, and the antiviral role of *Wolbachia*^5–10^. However, from an evolutionary perspective, the value of *D. melanogaster* is not just in its experimental tractability, but also in its close relationship to many other experimentally tractable species^11^. For example, experimental infection studies of more than 50 species of Drosophilidae (representing around 50 million years of evolution) have shown that susceptibility to viral infection has a strong phylogenetic component, such that more closely-related host species display more similar viral replication rates and virulence^12^, and that closer relatives of the virus’ natural host tend to support higher viral replication rates^13^. To understand how such phylogenetic patterns relate to host and virus biology in the wild we need to know the natural host range and frequency of host switching of these viruses. Thus, to capitalise on the value of the Drosophilidae as a model clade, we require a broader perspective on *Drosophila* viruses than *D. melanogaster* alone.

Prior to the advent of modern molecular biology, a handful of *Drosophila* viruses had been described on the basis of traditional virological techniques^14^. Starting with the Sigmavirus of *D. melanogaster* (DMelSV, Rhabdoviridae; shown to be a Rhabdovirus by ref. 15), which was initially identified by the failure of infected flies to recover from CO_2_ anaesthesia^16,17^, these ‘classical’ *Drosophila* viruses also include Drosophila P virus (DPV, Picornavirales^18^), *Drosophila C virus* (DCV, *Cripavirus*^19^), Drosophila A virus (DAV^20^), Drosophila F virus (Reoviridae^21^), and Drosophila G virus (Unclassified^21^) from adult flies, and *Drosophila X virus* (DXV, *Entomobirnavirus*^22^), Drosophila K virus (Reoviridae,^23^), and unnamed Reoviruses from cell culture (e.g. ref. 24, and see also ref 25). In broadly the same period, Iota virus (Picornavirales^26^) was identified from *D. immigrans* and was shown to be serologically similar to DPV, RS virus was identified in *D. ananassae* and members of the *D. montium* group^21^ and shown to be morphologically similar to Chronic Bee Paralysis virus, and Drosophila S virus (Reoviridae^27^) was identified from *D. simulans*. Unfortunately, of these ‘classical’ viruses, only DAV, DCV, DXV, and DMelSV remained in culture into the era of routine sequencing, and the others have been lost—making their classification tentative and relationships to each other and subsequently discovered viruses uncertain.

As large-scale sequencing became routine, it led to the serendipitous discovery of *Drosophila* viruses in host RNA sequenced for other purposes. Starting with the discovery of Nora virus (unclassified Picornavirales) in a *D. melanogaster* cDNA library^28^, such discoveries have included six viruses from small RNAs of *D. melanogaster* cell culture and *D. melanogaster* laboratory stocks (American Nodavirus, *D. melanogaster* totivirus, *D. melanogaster* Birnavirus, and Drosophila tetravirus^29^; *Drosophila* uncharacterized virus and *Drosophila* reovirus^30^), a novel Cripavirus in *D. kikkawai*^31^ and a new *Sigmavirus* in *D. montana*^32^. At the same time, PCR surveys of other *Drosophila* species using primers designed to *D. melanogaster* viruses were used to detect novel Nora viruses in *D. immigrans*, and *D. subobscura*^33^, and new Sigmaviruses in CO2-sensitive individuals of *D. affinis* and *D. obscura*^34^, and subsequently in *D. immigrans*, *D. tristis*, and *D. ananassae*^35^.

With the widespread adoption of high-throughput sequencing technologies the metagenomic (transcriptomic) sequencing of wild-collected flies is now starting to revolutionise our understanding of the drosophilid virome. The first explicitly metagenomic virus study in *Drosophila* discovered the first DNA virus of a drosophilid, *D. innubila* Nudivirus^36^. Subsequently, RNA and small-RNA sequencing of around 3000 *D. melanogaster* from the United Kingdom and 2000 individuals of several species from Kenya and the USA (primarily D. *melanogaster*, D. *ananassae, D. malerkotliana* and *Scaptodrosophila latifasciaeformis)* was used to identify more than 20 new RNA virus genomes and genome fragments, and a single near-complete DNA virus (Kallithea virus, Nudivirus)^31^. Metagenomic sequencing targeted to CO_2_ sensitive individuals has also recently been used to identify novel Sigmaviruses and other Rhabdoviruses in *D. algonquin*, *D. sturtevanti*, *D. busckii*, *D. subobscura*, *D. unispina*, and *S. deflexa*^32^.

In total, studies using classical virology, serendipitous transcriptomic discovery, and metagenomic sequencing have reported more than 60 viruses associated with the Drosophilidae and *Drosophila* cell culture (for a comprehensive list, see supporting information file 1). And, while the lost ‘classical’ viruses and incomplete metagenomic genomes make the exact number of distinct viruses uncertain, around 50 are currently represented by sequence data in public databases. From these it is possible to draw some general observations about the virus community of the Drosophilidae. For example, it is clear that RNA viruses substantially outnumber DNA viruses: of the *ca*. 50 viruses with published sequence, only two are DNA viruses (the Nudiviruses of *D. innubila*^36^ and *D. melanogaster*^31^). However, the extreme sampling bias introduced by targeted virus discovery, such as CO2-sensitivity analysis for Sigmaviruses (Rhabdoviridae^32^), makes it difficult to draw robust conclusions about the taxonomic composition of the *Drosophila* viruses. For example, among RNA viruses generally positive sense single stranded (+ssRNA) viruses are more common than other groups, but negative sense viruses (-ssRNA) constitute around 30% of classifiable *Drosophila* RNA viruses, and double-stranded (dsRNA) viruses nearly as high a proportion (supporting online file 1). To generalise such patterns, and to gain broader insight into the host-range of *Drosophila* viruses and their relationship to the viruses of other organisms, will require further unbiased metagenomic sequencing.

Here we report the viruses we have discovered through metagenomic sequencing of RNA from around 1600 wild-collected flies of the species *D. immigrans, D. obscura, D. subobscura, D. subsilvestris, D. tristis* and *S. deflexa*. We also report the re-analysis of two putatively virus-like sequences previously identified in a large pool of mixed *Drosophila*^31^. In total we describe 25 new viruses, and place these within the phylogenetic diversity of known viruses and undescribed virus-like sequences from public transcriptomic datasets. Remarkably, in wild *D. immigrans* we identify new viruses that are extremely closely related to the laboratory models DXV (previously known only from *Drosophila melanogaster* cell culture) and *Flock House virus* (originally isolated from beetles), and we detect the presence of *Armadillidium vulgare* iridescent virus^37^ in *D. immigrans* and *D. obscura*—only the third DNA virus to be reported in a drosophilid. We find that a few viruses, such as La Jolla virus^31^, appear to be generalists, and that many viruses are shared between the closely-related members of the *Drosophila obscura* group, but that viruses are more rarely shared between more distantly-related species. We discuss our findings in the context of the Drosophilidae as a model clade for studying host-virus coevolution, and the diversity and host range of invertebrate viruses more generally.

## Methods

### Sample collections and sequencing

We collected around 1400 adult flies representing five species in the United Kingdom in summer 2011 (*D. immigrans, D. obscura, D. subobscura, D. subsilvestris*, and *D. tristis*) and 200 *Scaptodrosophila deflexa* in France in summer 2012. Flies were netted or aspirated from banana/yeast bait in wooded and rural areas at intervals of 24 hours for up to a week at each location. They were sorted morphologically by species, and RNA was extracted using Trizol (Ambion) according to the manufacturer’s instructions. Females of the obscura group (including *D. obscura, D. subobscura, D. subsilvestris*, and *D. tristis*) are hard to identify morphologically, and for these species only males were used for RNA extraction and sequencing.

In total, 498 *D. immigrans* were collected in three groups (63 flies in July 2011 Edinburgh 55.928N, 3.170W; 285 flies in July 2011 Edinburgh 55.921N, 3.193W; 150 flies July 2011 Sussex 51.100N, 0.164E). The 502 *D. obscura* males were collected in four groups (280 flies collected in July 2011 Edinburgh 55.928N, 3.170W; 52 flies October 2011 Edinburgh 55.928N, 3.170W; 115 flies July 2011 Sussex 51.100N, 0.164E; 55 flies August 2011 Perthshire 56.316N, 3.790W). The 338 *D. subobscura* males were collected in four groups (6g flies collected in July 2011 Edinburgh 55.928N, 3.170W; 60 flies in October 2011 Edinburgh 55.928N, 3.170W; 38 flies July 2011 Sussex 51.100N, 0.164E; 180 flies August 2011 Perthshire 56.316N, 3.790W). The 64 *D. subsilvestris* were collected in three groups (44 flies collected in July 2011 Edinburgh 55.928N, 3.19W; 15 flies in October 2011 Edinburgh 55.928N, 3.19W; 5 flies in August 2011 Perthshire 56.316N, 3.790W). The 29 *D. tristis* were collected in two groups from a single location (21 flies collected July 2011 Edinburgh 55.928N, 3.190W; 8 flies October 2011 Edinburgh 55.928N, 3.190W), and the approximately 200 *S. deflexa* in a single collection (August 2012 in Le Gorges du Chambon, France 45.66N, 0.556E). Pooled cytochrome oxidase sequence data subsequently showed that some of these collections may be contaminated with other species. Specifically, around 2% of reads in the *D. subobscura* sample appear to derive from *D. tristis*, and around 5% of reads in the *D. subsilvestris* sample may derive from *D. bifasciata*.

RNA was treated with DNAse (Turbo DNA-free, Ambion) to reduce DNA contamination, and precipitated in RNAstable (Biomatrica) for shipping. All library preparation and sequencing was performed by the Beijing Genomics Institute (BGI tech solutions, Hong Kong) using the Illumina platform and either 91nt or 101nt paired-end reads. Raw data are available from the sequencing read archive (SRA) under project accession SRP070549. Initially, two separate sequencing libraries were prepared for *D. immigrans*, the first used Ribo-Zero (Illumina) depletion of rRNA to increase the representation of viruses and host mRNAs (SRR3178477), and the second used duplex-specific nuclease normalisation (DSN) to increase the representation of rare transcripts (SRR3178468). Subsequently, for each of the other species a single library was prepared, again using DSN normalisation (*D. obscura* SRR3178507; *D. subobscura* SRR3180643; *D. subsilvestris* SRR3180644; *D. tristis* SRR3180646; *S. deflexa* SRR3180647). Unfortunately, due to a miscommunication with the sequencing provider, these six libraries were subject to poly-A selection prior to normalisation. This process substantially increases the amount of virus sequence available for assembly and identification (by excluding rRNA), but will bias viral discovery toward virus genomes and sub-genomic products that are poly-adenylated (e.g. Picornavirales). Sequencing resulted in an average of 48 million read pairs per library, ranging from 47.3M read pairs for *D. subobscura* to 52.7M read pairs for the *D. immigrans* DSN library.

### Virus genome assembly and identification

Raw reads were quality-trimmed using sickle (version 1.2^38^) only retaining reads longer than 40nt, and adapter sequences were removed using cutadapt (version 1.8.1^39^). Paired-end sequences were then *de novo* assembled using Trinity (version 2.0.6^40^) with default parameters, and the resulting raw unannotated assemblies are provided in supporting information file 2. In the absence of confirmation (e.g. by PCR) such assemblies necessarily remain tentative, and may represent chimeras of related sequences or contain substantial assembly errors.

We took two approaches to identify candidate ‘virus-like’ contigs for further analysis. First, for each nominal gene assembled by Trinity, we identified and translated the longest open reading frame, and used these translations to query virus sequences present in the Genbank non-redundant protein database (‘nr’)^41^ using blastp (blast version 2.2.28+)^42^ with default parameters and an e-value threshold of 10^−5^, and retaining the single ‘best’ hit. Second, for each nominal gene, we used the transcript with the longest open reading frame to query virus sequences in ‘nr’ using blastx with default parameters, but again using an e-value threshold of 10^−5^ and retaining the single best hit. These two candidate lists, comprising all the sequences for which the top hit was a virus, were then combined and used to query the whole of nr using blastp, using an e-value threshold of 10^−5^ and retaining the top 20 hits. Sequences for which the top hit was still a virus, and sequences with a blastx hit to viruses but no other blastp hits in nr, were then treated as putatively viral in origin, and subject to further analysis. In parallel with these analyses, raw data that we previously reported from *D. melanogaster*^31^ were re-assembled and re-analysed in the same way.

For each putative virus fragment we selected other virus-like fragments in the same host that showed sequence similarity to the same virus taxonomic group, e.g. combining all Negevirus-like sequences in *D. immigrans*, or all Rhabdovirus-like sequences in *D. obscura*. We then manually ordered and orientated these fragments by reference to the closest relatives in Genbank to identify longer contigs that had not been assembled by Trinity. In some cases we were able to identify very long contigs (i.e. near-complete viral genomes) in the Genbank Transcriptome Shotgun Assembly database (‘tsa_nt’), and use these to order, orientate, and join overlapping virus fragments that had remained un-joined in the Trinity assembly. In cases of ambiguity, for example where fragments failed to overlap and related viruses were present in the same pool, we did not manually join contigs. Where helpful, we used the longer TSA sequences to query our *Drosophila* metagenomic data using tblastx and thereby identify further fragments to complete viral genomes. Near-complete genome sequences from Nora viruses of *D. immigrans* and *D. subobscura*, and Sigmaviruses of *D. tristis* and *S. deflexa*, were reported previously, and are not further analysed here^32,33^. The remaining novel virus contigs are reported here and have been submitted to Genbank under accession numbers KU754504-KU754539.

### Re-analysis of RNA data from D. melanogaster

Blast analysis suggests that two of the putative viral genomes identified during the course of this study (Hermitage virus of *D. immigrans*, and Buckhurst virus of *D. obscura;* see Results) are close relatives of short virus-like contigs that had previously been identified in *D. melanogaster* (previous contigs available from doi:10.1371/journal.pbio.1002210.s002; ref. 31). We therefore used the new longer contigs from *D. immigrans* and *D. obscura* to guide the assembly of (partial) genomes for the *D. melogaster* viruses. As small RNA data were available for the published *D. melanogaster* samples (data available user the SRA accession SRP056120; ref 31) we additionally mapped small RNAs to these viral genomes using Bowtie2^43^ to examine their properties.

### Phylogenetic analysis

We inferred the phylogenetic placement of each virus using a conserved region of coding sequence. Where possible, this was the RNA polymerase, as these tend to be highly conserved in RNA viruses. We used blastp to query the Genbank non-redundant protein database (nr) and tblastn to query the Genbank Transcriptome Shotgun Assembly database (tsa_nt) to identify potential relatives for inclusion in the phylogenetic analysis. For viruses that could be tentatively assigned by blast to a well-studied group (e.g. Iflaviruses, Nodaviruses), we additionally selected key representative members of the clade from the NCBI Virus genomes reference database^44^. We aligned protein sequences using Mcoffee from the T-Coffee package^45^, combining a consensus of alignments from ClustalW^46^, T-coffee^45^, POA^47^, Muscle^48^, Mafft^49^, DIALIGN^50^, PCMA^51^ and Probcons^52^. Consensus alignments were examined by eye, and the most ambiguous regions of alignment at either end removed. Nevertheless, as expected for an analysis of distantly-related and rapidly-evolving RNA viruses, these alignments retain substantial ambiguity and more distant relationships within the resulting phylogenetic trees should be treated with caution. Alignments are provided in supporting information file 3.

Alignments were used to infer maximum-likelihood trees using PhyML (version 20120412)^53^ with the LG substitution model^54^, empirical amino-acid frequencies, and a four-category gamma distribution of rates with an inferred shape parameter. Maximum parsimony trees were used to provide the starting tree for the topology search, and the preferred tree was the one with the highest likelihood identified after both nearest-neighbour interchange (NNI) and sub-tree prune and re-graft (SPR) searches. Support was assessed in two ways, first using the Shimodaira-Hasegawa-like nonparametric version of an approximate likelihood ratio test (see ref. 55) as implemented in PhyML, and second by examining 100 bootstrap replicates.

### Origin of RNA sequence reads

To infer the proportion of reads mapping to each virus, and to detect potential cross-species contamination in the fly collections, quality-trimmed reads were mapped to all the new and previously published drosophilid virus genomes, and to a 343 nt region of Cytochrome Oxidase I that provides a high level of discrimination between drosophilid species. Mapping was performed using Bowtie 2 (version 2.2.5)^43^ with default parameters and global mapping, and only the forward read in each read pair was mapped. To reduce the potential for cross-mapping between closely-related sequences, we excluded all trimmed reads with fewer than 80 contiguous non-N characters.

## Results

In total we identified 25 new RNA viruses through metagenomic sequencing of wild caught Drosophilidae. Among those viruses that could easily be classified were four members of the Picornavirales, three Rhabdoviruses, two Nodaviruses, two Reoviruses and an Entomobirnavirus (Fig 1). Among those lacking a current classification were five viruses distantly related to Negeviruses, four viruses distantly related to Sobemoviruses and Polerovirus, two distantly related to Flaviviruses, and two distantly related to Tombusviruses (Fig 2). It is striking that among this latter group there are many viruses that are closely related to unrecognised virus-like sequences in transcriptomic data. Indeed, of the 355 sequences we included in our phylogenetic analyses, nearly one third (29%) were derived from transcriptome data rather than from published viruses, illustrating the under-sampling of RNA viruses generally. All phylogenetic trees, including node-support values and Genbank accession numbers, are provided in supporting information file 4.

**Figure 1:**
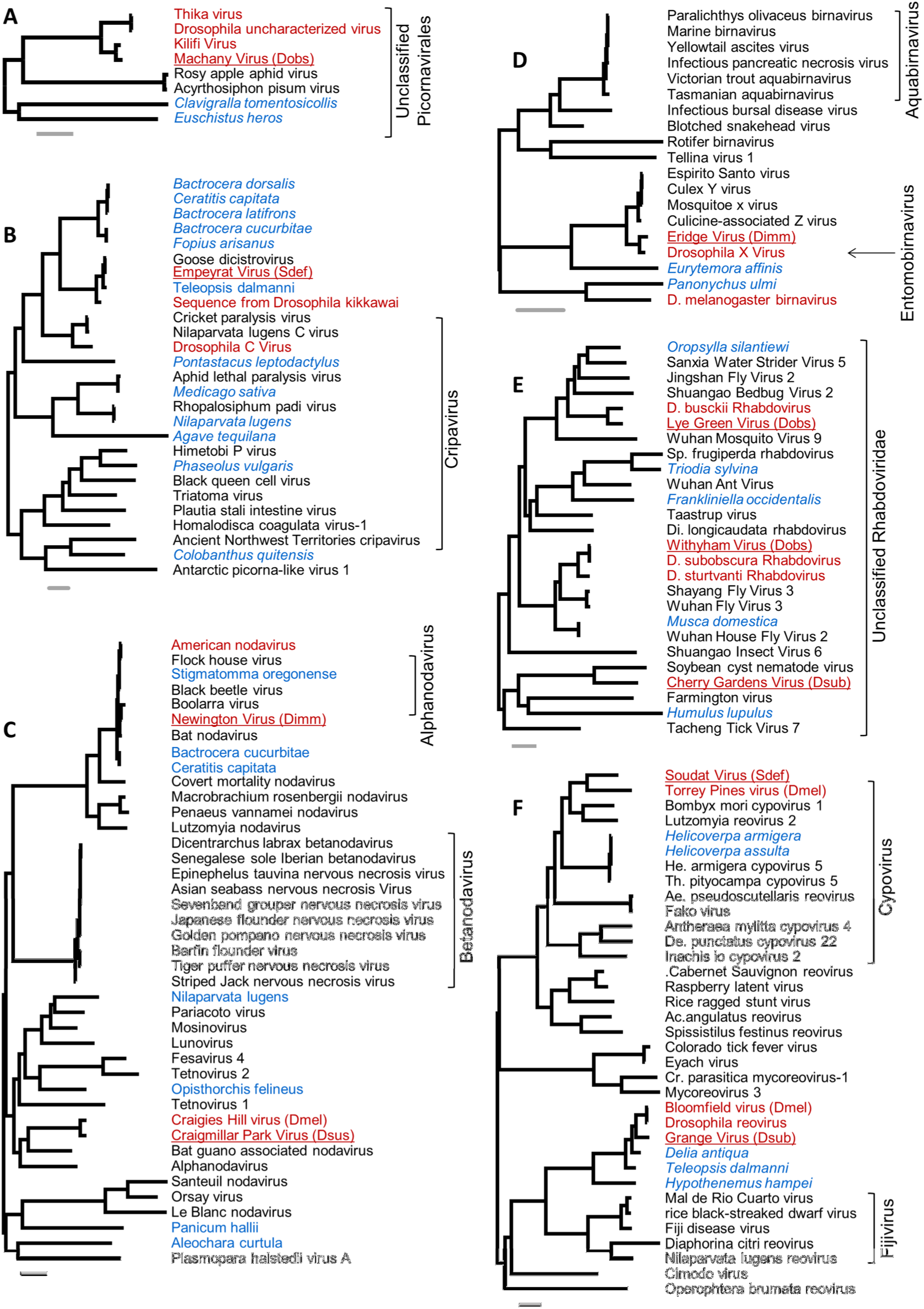
Viruses related to well-studied clades. Mid-point rooted maximum-likelihood phylogenetic trees for the viruses reported here, inferred using polymerase protein sequences. The grey scale bars represent 0.5 amino-acid substitutions per site. In each tree, viruses reported from Drosophilidae in labelled in red, viruses from other taxa are labelled in black, and unannotated virus-like sequences from publicly-available transcriptome datasets labelled in blue. Viruses newly-reported here are underlined, and *Drosophila* species abbreviations are given for the reference sequence (Dimm – *D. immigrans*; Dobs – *D. obscura*; Dsub – *D. subobscura*; Dsus – *D. subsilvestris;* Dtri – *D. tristis;* Sdef – *Scaptodrosophila deflexa*). Tree A: Viruses near to the Dicistroviridae (Picornavirales). B: Putative Cripaviruses (Dicistroviridae, Picornavirales – the corresponding tree in Supporting File 4 additionally includes Aparaviruses). C: Nodaviruses. D: Birnaviruses. E: Unclassified members of the Rhabdoviridae that form the sister clade to the Cytorhabdoviruses and the Nucleorhabdoviruses^32^. F: Reoviridae. Alignments are provided in online supporting file 3, and clade support values and sequence accession identifiers are provided in online supporting file 4.

**Figure 2:**
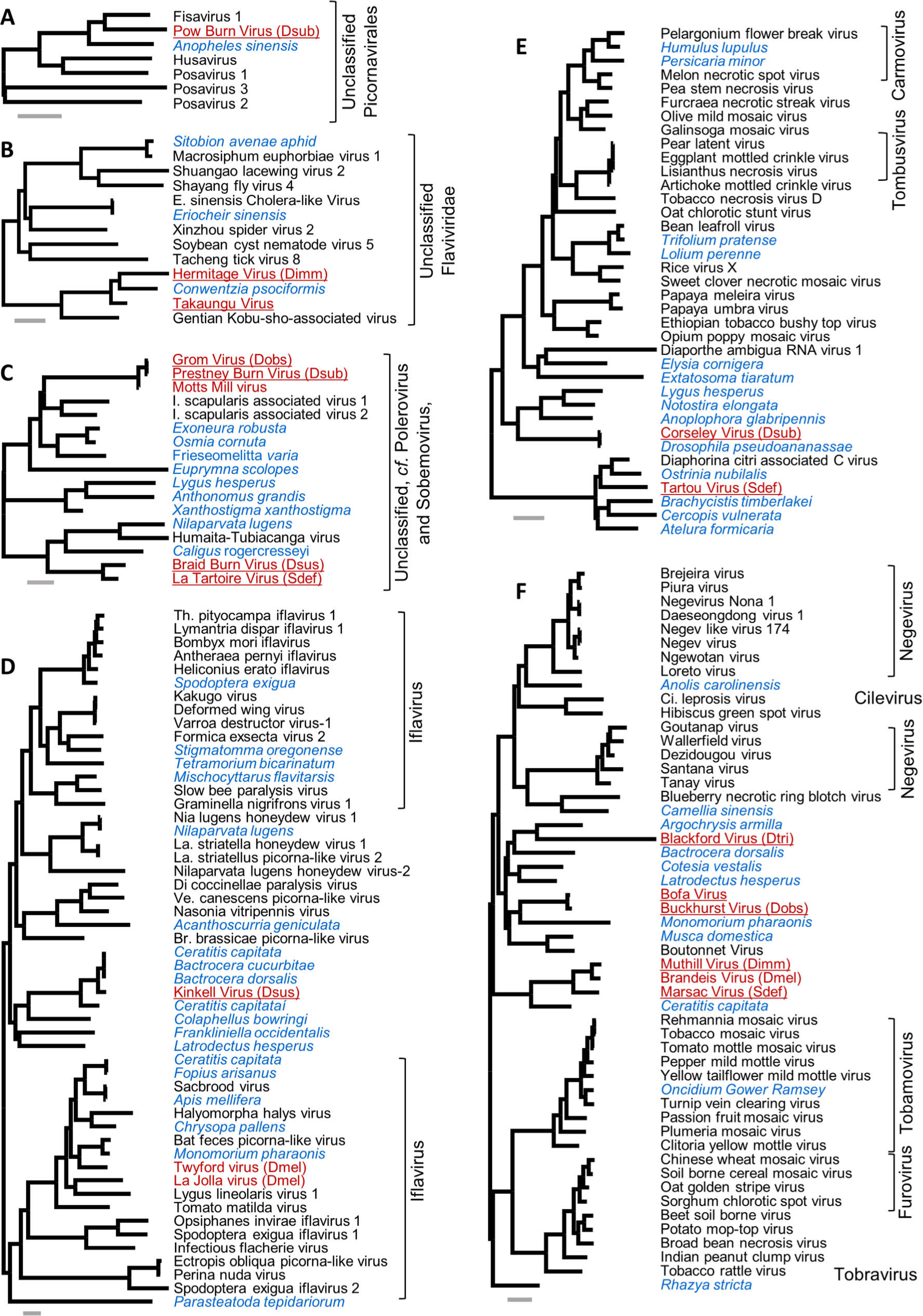
Viruses not closely related to well-studied clades. See Fig. 1 for a key to the colours and abbreviations. Tree A: Unclassified Picornavirales. B: Unclassified clade of basally-branching Flavi-like viruses^75^. C: An unclassified clade that branches basally to Poleroviruses and Sobemoviruses^79^. C: Nodaviruses. D: Iflaviruses, including a new clade that falls within (or close to) the Iflaviruses. E: Two unclassified clades related to the Tombusviridae^73^. F: Two unclassified clades related to the Negeviruses and the Virgaviridae. Alignments are provided in online supporting file 3, and clade support values and sequence accession identifiers are provided in online supporting file 4.

Following common practice, we have provisionally named the new *Drosophila* viruses after localities near to our collection sites. We have chosen this approach as it avoids associating the sequence with higher levels of either the host or virus taxonomy, when both may be uncertain or unstable. The new *Drosophila* viruses are each represented by between 1.8 kbp and 13.7 kbp of sequence (Tartou virus of *S. deflexa*, and Lye Green virus of *D. obscura*, respectively), and six are likely to be near-complete genomes with more than 9 kbp of sequence each. We have not named, and do not report, virus sequences that were near-identical to previously published viruses (i.e. *K*_*s*_<0.3, or falling within the published diversity of other viruses). See Fig 3 for read numbers of previously published viruses.

**Figure 3:**
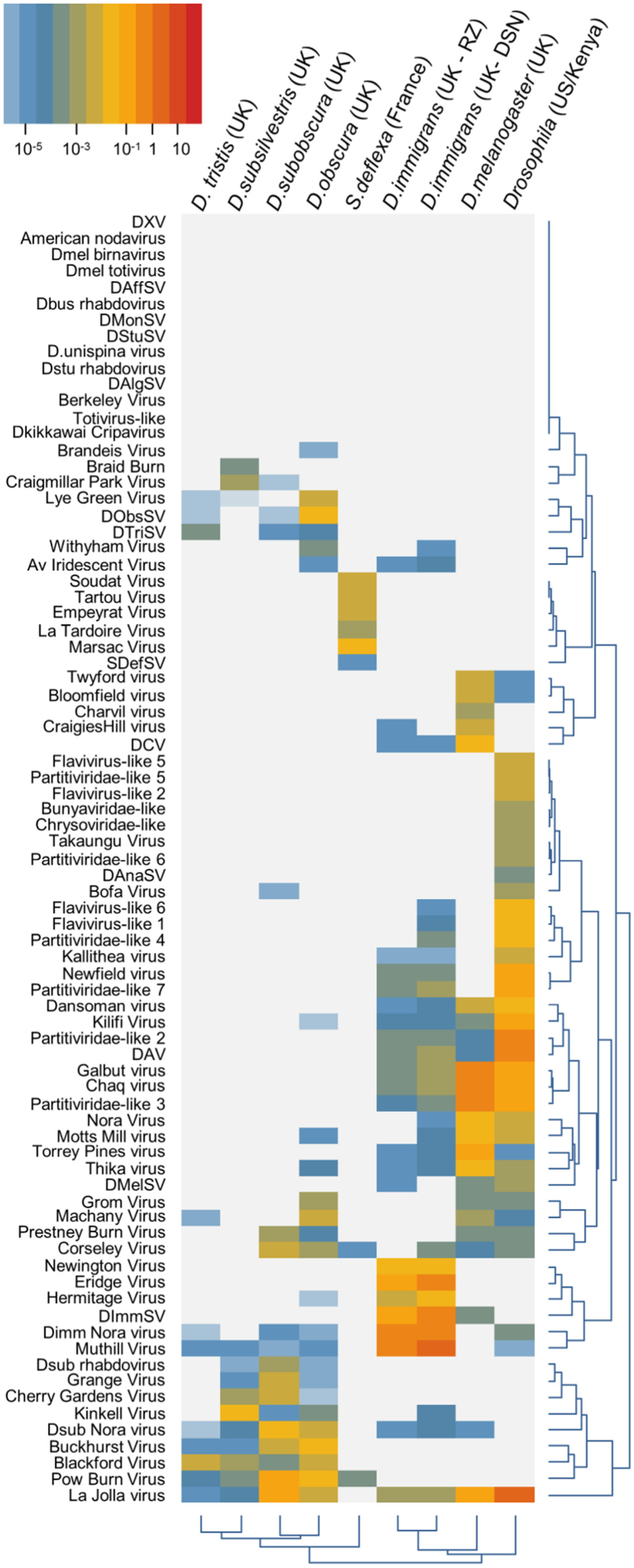
Virus read numbers. A heatmap showing the relative number of high quality (8G nt) forward reads from each library that map to each of the *Drosophila* viruses. Read numbers are normalised by target sequence length, and by the number of reads mapping to a fragment of the host Cytochrome Oxidase 1 gene (so that a value of 1 implies equal read numbers per unit length of the virus and the host Cytochrome Oxidase I). Rows and columns are clustered by the similarity in read frequency on a log scale. Note that some viruses may be sufficiently similar for a small proportion of reads to cross-map, and that a small level of cross-contamination between fly species means that the data presented here cannot be used to confidently infer host-specificity.

### New viruses closely related to viruses of D. melanogaster

For around half of the newly-discovered viruses (11 of 25), the closest previously-reported relative was associated with *D. melanogaster*. Most striking of these is Eridge virus, a segmented dsRNA Entomobirnavirus closely related to the *D. melanogaster* laboratory model, *Drosophila X virus*^56^ (Fig 1 D; 78% sequence identity and 83% amino-acid identity in Segment A). DXV has not previously been observed in wild flies, but has been reported from flies injected with fetal bovine serum and has therefore been considered a cell culture contaminant^57^. In addition to DXV, we detected sequences that were >98% identical to Eridge virus in some *Drosophila* cell cultures (e.g. ModEncode dataset SRR1197282 from S2-DRSC cells^58^), showing that fly cell cultures can harbour both viruses.

Other viruses that are also closely related to a published *Drosophila* virus include Machany virus of *D. obscura* (unclassified Picornavirales, close to Kilifi virus and Thika virus of *D. melanogaster;* Fig 1 A), Grange virus of *D. subobscura* (a Reovirus close to Bloomfield virus of *D. melanogaster;* Fig 1 F), Craigmillar Park virus of *D. subsilvestris* (an Alphanodavirus close to Craigie’s Hill virus of *D. melanogaster;* Fig 1 C), Grom virus and Prestney Burn virus (of *D. obscura* and *D. subobscura* respectively, both close to Motts Mill virus of *D. melanogaster*; Fig 2 C), and Muthill virus and Marsac virus (of *D. immigrans* and *S. deflexa* respectively Fig 2 F; both close to Brandeis virus identified in publicly available *D. melanogaster* sequence data from laboratory stocks^31,59^).

### New Drosophila viruses closely related to viruses of other species

We identified two new viruses that are extremely closely related to viruses reported from other taxa. Newington virus of *D. immigrans* is an Alphanodavirus extremely similar to *Boolarra virus*^60^ (isolated from the lepidopteran *Oncopera intricoides;* 84% nucleic acid identity and 89% amino-acid identity in the polymerase), the widely-used laboratory model *Flock House virus*^60^ (from the coleopteran *Costelytra zealandica*; 79% nucleic acid and 87% amino acid identity) and American Noda virus (ANV, identified from small RNAs of *D. melanogaster* cell culture^29^). This clade of closely-related nodaviruses also includes Bat Nodavirus (detected in the brain tissue of the insectivorous bat *Eptesicus serotinus*^61^) and transcriptome sequences from the flies *Bactrocera cucurbitae*^62^ and *Ceratitis capitate*^63^.

We further identified a novel Cripavirus in *S. deflexa* that is very closely related to Goose Dicistrovirus (90% sequence identity, 92% amino-acid identity), recently identified from a faecal sample from geese^64^. However, given that the next closest relatives to this sequence are a transcriptome sequence from the stalk-eyed fly *Teleopsis dalmanni*^65^, and a Cripavirus present in publicly available transcriptome data from *D. kikkawai* (supplementary information in ref. 31), we think it likely that these represent invertebrate viruses. To reflect this, and given the divergence between them, we have decided to consider the *S. deflexa-*associated sequence as a new virus, and have provisionally named it Empeyrat virus.

### New viruses without close relatives

The remaining other new putative viruses (13 of 25) do not have published close relatives, although many are related to unreported viruses present in host transcriptome datasets. Most notable among these are Kinkell virus of *D. subsilvestris* and Corseley virus of *D. subobscura*. Kinkell virus, along with transcriptome sequences from the fly genera *Bactrocera*^62,66^ and *Ceratitis*^63^, the beetle *Colaphellus*^67^, the thrip *Frankliniella*^68^, and the spider *Latrodectus*^69^, appears to define a major new clade that falls within or close to the Iflaviruses (Fig 2 D). Similarly, Corseley virus, which is almost identical to transcriptome sequences from *D. pseudoananassae*^70^ and is related to transcriptome sequences from the bug genus *Lygus*^71^ and the beetle genus *Anoplophora*^72^, appears to define an entirely new group of viruses distantly related to Tombusviridae and the recently-described *Diaphorina citri* associated C virus^73^ (which is itself closely related to the newly identified Tartou virus of *S. deflexa;* Fig 2 E).

Two other groups are also noteworthy. First, the clade that includes Takaungu virus—which we have identified through re-analyses of mixed Drosophilid sequences from Kenya^31^—and Hermitage virus of *D. immigrans*. These viruses are most closely related to a transcriptome sequence from the Neuropteran *Conwentzia psociformis*, and the enigmatic Gentian Kobu-sho-associated virus, which is reported to be an extremely large dsRNA relative of the Flaviviruses (Fig 1 B; ref. 74, see also ref. 75). Second, is the clade that includes Blackford virus of *D. tristis*, Buckhurst virus of *D. obscura*, and Bofa virus (also derived from the Kenyan pool^31^, incorporating three unnamed fragments KP757936, KP757935, and KP757975). These viruses, along with seven transcriptome sequences from various arthropods and Muthill, Marsac, and Bradeis viruses (described above), appear to represent a major group of insect-infecting viruses that fall between the recently proposed Negeviruses^76^ and the plant virus family Virgaviridae.

### A DNA Iridescent virus in Drosophila

In *D. immigrans* and *D. obscura* we identified more than 900 read pairs almost identical to the DNA iridescent virus of *Armadillidium vulgare* (Invertebrate Iridovirus 31, ref. 37). Although read numbers were relatively small (around 700 high-quality read pairs in *D. obscura* and 250 read pairs in *D. immigrans*), they do not represent low-complexity sequence, they are widely distributed around the viral genome, and they suggest that viral genes were being expressed (i.e. present in RNA). The longest contiguous region of coverage in *D. obscura* corresponded to the virus major capsid protein, and displayed 98% sequence identity to *Armadillidium vulgare* DNA iridescent virus (*K*_*s*_=0.08). These data suggest that this virus has a broad host range, and represent the third DNA virus to be identified naturally infecting a drosophilid.

### Small RNA data from Takaungu virus and Bofa virus

For Takaungu virus (Contigs KP757925 and KU754513) and Bofa virus (KU754515) small (19-30nt) RNA data were available from our previous study of *D. melanogaster*^31^. Although relatively few small RNA reads were detected from these viruses (*ca*. 200 reads from Bofa virus, *ca*. 800 reads from Takaungu virus), the small RNAs displayed the properties expected of virus-derived siRNAs in *Drosophila* (Supporting online file 5). Specifically, they derived from both strands of the virus, they were distributed along the full length of the virus contigs, their size distribution peaked sharply at 21nt (in contrast to viral siRNAs of chelicerates, hymenopterans, and nematodes that are predominantly 22nt in length), and there was a bias against G in the 5' position.

### The distribution of virus reads across host species

To explore the distribution of viruses across hosts, we mapped high-quality reads from all libraries to new and previously reported *Drosophila* virus sequences (Fig. 3). We included a UK sample of *D. melanogaster* and a mixed drosophilid pool from Kenya and the USA that we published previously^31^. Overall, approximately 1% of RNAseq reads were viral in origin, ranging from 0.02% in the *D. tristis* pool to 6.96% in the mixed drosophilid pool. As expected, many published *Drosophila* viruses were absent. These include all the Rhabdoviruses from host species not present in our collections (Rhabdoviruses from *D. affinis, D. busckii, D. montana, D. surtevanti, D. algonquin* and *D. unispina*,) and the Cripavirus identified in public RNA reads from *D. kikkawai*^31^ (host also absent from our collections). Absent viruses also included the five that have previously only been identified in cell culture (*Drosophila X virus*, American Nodavirus, *D. melanogaster* Birnavirus, *D. melanogaster* Totivirus^29^, and the totivirus from public dataset SRR1197466 ^31^), and also Berkeley virus (identified in reads from SRR070416^31^).

The number of viruses varied substantially among metagenomic pools. Normalising by sequence length and by the number of reads from host Cytochrome Oxidase 1 (to account for variation in total read numbers, rRNA contamination levels, and sequence lengths) we were able to detect between 4 viruses (*D.tristis*) and 27 viruses (*D. immigrans* DSN) per pool at 0.001% of COI expression. The number of detectable viruses was positively correlated with the number of flies in the single-species samples, and the strength of the relationship increased with the expression threshold for inclusion (Spearman rank correlations: at 0.001% of COI ρ=0.86, *p*=0.02; at 0.01% of COI ρ=0.96, *p*=0.0008; and at 0.1% of COI ρ=0.96, *p*=0.003). For *D. immigrans*, the DSN library detected more viruses than the rRNA depleted library, regardless of threshold. Note that the presence of some cross-mapping between related viruses means that estimates of the number of viruses will tend to be slightly inflated at low thresholds.

Although our sampling scheme and a small amount of species cross-contamination precludes a rigorous formal analysis of host range, some viruses do appear to be generalists and others specialists. Using the 0.01% threshold, the majority of Rhabdoviruses (including Sigmaviruses) appeared to be restricted to a single host: assuming the apparent low level of DImmSV in *D. melanogaster* is due to cross-mapping, only Cherry Gardens virus (related to Soybean cyst nematode associated northern cereal mosaic virus^77^) was present in two host species (*D. subobscura* and *D. subsilvestris*). In contrast, a few viruses appeared to have a broad host range: La Jolla virus (Iflavirus), Blackford virus (related to Negeviruses and the Virgaviridae), Corseley virus (related to Tombusviruses), and Pow Burn virus (Picornavirales, related to Fisavirus 1) were each present in four species at >0.01% COI, and small numbers of La Jolla virus reads were detected in all pools except *S. deflexa*. Considering read frequencies across all viruses, members of the obscura group displayed the greatest similarity to each other (Fig.3; *D.obscura, D. subobscura, D. tristis* and *D. subsilvestris*), while *S. deflexa* was the most distinct, with 6 of its viruses not present in any other pool, and only two of the viruses from other pools present in *S. deflexa*.

## Discussion

### New viruses of Drosophila

The twenty five new viruses we present here bring the total number of viruses reported from the Drosophilidae to approximately 85 (see supporting information file 1). However, although it does not detract from the potential utility of the viruses we were able to identify, it should be noted that this sampling is far from comprehensive. First, more viruses are likely to have been present in these samples than we were able to detect—for example, because viral titre was too low for some viruses or (for flies other than *D. immigrans*) because poly-A selection biases against their discovery. Second, more virulent viruses may reduce fly movement, so that virulent viruses are underrepresented by collections from baited traps.

As for the majority of metagenomic studies, it also remains uncertain whether these viruses constitute active infections of *Drosophila*, or whether they are contaminants of the host surface or gut lumen, infections of an unrecognised parasite or other Drosophila-associated microflora, or ‘fossil’ endogenous viral elements integrated into the host genome and still expressed (‘EVEs’^78^). Small RNA sequencing can in principle be used to demonstrate that viruses do replicate within arthropod and nematode hosts, and are targeted by their immune system^30,31^. In addition, as hymenoptera, chelicerata, and nematodes generate predominantly 22nt small RNAs from viruses, the presence of 21 nt virus-derived siRNAs is highly suggestive of an immune response by *Drosophila*. As two of the viruses reported here (Takaungu virus and Bofa virus) were identified through re-analysis of data from Webster *et al*^31^, we were able to test whether these viruses show the expected siRNA profile. As expected, we do detect 21nt siRNAs from both strands of these two viruses, consistent with their replication in *Drosophila* (supporting online file 5). Indeed, in the earlier analysis^31^ we identified an unnamed but putatively viral sequence purely on the basis of 21nt siRNAs that can now be shown to be part of Bofa virus (Genbank accession KP757975; sufficient similarity to identify Bofa virus using blast is now provided by Buckhurst virus).

Nevertheless, in the absence of small RNA data for the other 23 putative viruses presented here, it remains possible that these virus-like sequences are EVEs^78^, or infections o*f Drosophila*-associated microflora. However, while EVEs are common in insect genomes^78^, expressed EVEs are rarer, and expressed EVEs appear to be extremely rare relative to active viral infections. For example, in our previous metagenomic study of *Drosophila* RNA viruses, none of the 14 viruses we initially identified by RNA sequencing in *D. melanogaster* proved to be EVEs^31^. Thus, although a minority of the sequences presented here could be recently acquired EVEs, few are likely to be as they do not appear in the genomes of closely related hosts, they are expressed, and they appear to be constrained (we detect long open reading frames).

Fifteen of remaining 23 putative viruses in the present study are extremely closely related to known insect viruses or virus-like sequences from insect transcriptomes (Fig 1, Fig 2), and/or are present at such high levels (greater than 10% of host COI in the cases of Muthill virus and Eridge virus), that is seems likely that the associated drosophilid is indeed the host. For the remaining eight—namely Braid Burn virus, Cherry Gardens virus, Blackford virus, La Tardoire virus, Hermitage virus, Pow Burn virus, Tartou virus, and Soudat virus—conclusive demonstration of a drosophilid host must await future siRNA sequencing or experimental confirmation.

Three groups of newly discovered and currently unclassified viruses seem particularly prominent within the drosophilid samples presented here. First, near to the Sobemoviruses and Poleroviruses are a large clade of invertebrate-infecting viruses defined by Ixodes Tick Associated viruses 1 and 2 (ref 79), Humaita-Tubiacanga virus^30^, the Drosophila-associated Grom virus, Prestney Burn virus, Motts Mill virus, Braid Burn virus and La Tardoire virus, and transcriptome-derived sequences predominantly from Hymenoptera and Hemiptera. Second, branching basally to the Negeviruses (and potentially between Negeviruses and Virgaviridae) are two clades including the *Drosophila-associated* Blackford virus, Bofa virus, Buckhurst virus, Brandeis virus, Muthill virus and Marsac virus, along with transcriptome-derived sequences dominated by Diptera and Hymenoptera. Third, near to the Tombusviridae are the clades defined by *Diaphorina citri* associated C virus^73^, Tartou virus and Corseley virus from the Drosophilidae, and transcripts from various invertebrates. All three groups appear to represent common and widespread infections of invertebrates that warrant taxonomic recognition.

### Virus diversity and host range

Rapid viral discovery, facilitated by large-scale metagenomic sequencing and the serendipitous discovery of viral genomes in transcriptomic data, is revolutionising our understanding of virus diversity. The Drosophilidae provide a clear example of this, with approximately ten viruses reported prior to the year 2000, eleven more between 2001 and 2014, and more than 60 since 2015. Particularly striking is the frequency with which completely new, and deeply divergent, lineages of RNA viruses are being identified. Recent examples include the enormous and unexpected diversity of basally-branching ‐ssRNA viruses^80^, the diversity of basal Flaviviridae^75^, the Negeviruses^76^, and the Phasmaviruses^81^.

How many invertebrate viruses are there, and when will the accelerating virus-discovery curve start to saturate? Our *ad hoc* but intensive sampling of *Drosophila* suggests that such questions will require systematic estimates of the distribution of virus host ranges, the distribution of virus geographic ranges, and the distribution of virus prevalences. First, many *Drosophila* viruses are multi-host and widely-distributed. Around 10 of the 25 new viruses we report are detectable in multiple species, and we also detect previously published viruses of *D. melanogaster* in *D. immigrans* and members of the obscura group (Fig. 3). Similarly, our earlier PCR survey of *D. melanogaster* viruses^31^ detected 12 of 16 viruses in more than a third of *D. melanogaster* populations, and 10 of them in at least one *D. simulans* population. Second, it seems likely that more closely-related hosts share more viruses. This is consistent with the apparently high overlap in virus community between *D. melanogaster* and *D. simulans*^31^ and among members of the obscura group, and the divergent set of viruses associated with *S. deflexa* (Fig. 3, but note that the *D. subobscura* sample was slightly contaminated by *D. tristis*, and the *D. subsilvestris* sample by *D. bifasciata*). It is also consistent with the absence of *D. melanogaster* viruses from metagenomic surveys of other invertebrate taxa (although Goose dicistrovirus is closely related to Empeyrat virus of *S. deflexa*). Third, it is clear that viruses vary enormously in prevalence, such that few viruses are common and many are rare. Of the 16 viruses we previously surveyed by PCR, only three ever exceeded 50% prevalence, and most only exceeded 10% prevalence in two or three of the surveyed populations. This is consistent with the positive relationship we find between sample size and virus number, and suggests that many hundreds of *Drosophila* individuals are required to comprehensively survey a population.

## Conclusions

The 25 new viruses we present here expand the catalogue of recorded drosophilid-associated viruses by nearly 50%, and identify several new clades of insect-associated viruses. These include a new clade related to the Iflaviruses (Kinkell virus), new clades related to the Tombusviridae (Corseley virus and Tartou virus), and new clades related to the Negeviruses and Virgaviridae (including 6 viruses detected in *Drosophila*). Nevertheless, the large numbers of undescribed viruses present in transcriptome datasets illustrates that, across the invertebrates as a whole, there are many more viruses and many more deeply divergent virus lineages to uncover.

We expect that the future isolation of these Drosophila-associated viruses will provide useful laboratory tools to better understand host-virus biology and host range. However, it is possible to capitalise on viral sequences to address these questions even in the absence of viable viral isolates, and new virus sequences *per se* are likely to prove valuable^33^. In addition, given the widespread experimental use of model viruses that are not known to infect *D. melanogaster* in the wild, such as *Flock house virus*^82,83^, *Drosophila X virus*^84,85^, and Invertebrate Iridovirus ? (ref. 86), it is reassuring to know that these viruses have close relatives naturally associated with the Drosophilidae (respectively Newington virus, Eridge virus, and *Armadillidium vulgare* iridescent virus in *D. immigrans*).

## Acknowledgements

We thank the City of Edinburgh Council, Friends of the Hermitage of Braid, Keith and Sue Obbard, Sandy and Helen-May Bayne, Elizabeth Bayne, Graeme Pratt, and Neil and Jacky Longdon for permission to collect *Drosophila* and/or for support while making collections. We thank Fergal Waldron and Gytis Dudas for discussion.

## Funding sources

This work was funded by a Wellcome Trust Research Career Development Fellowship to DJO (WT085064). BL was supported by grants from the UK Natural Environment Research Council (NE/L004232/1) and the European Research Council (281668, DrosophilaInfection). SHL was supported by a Natural Environment Research Council Doctoral Training Grant (NERC DG NE/J500021/1). Work in DJO’s lab is partly supported by a Wellcome Trust strategic award to the Centre for Immunity, Infection and Evolution (WT095831).

## Author Contributions

Conceived and designed the experiments: DJO BL. Performed experiments: DJO BL SHL CLW. Analysed the data: DJO. Wrote the first draft of the manuscript: DJO. Contributed to the writing of the manuscript: DJO BL SHL CLW.

---

**Table 1:**
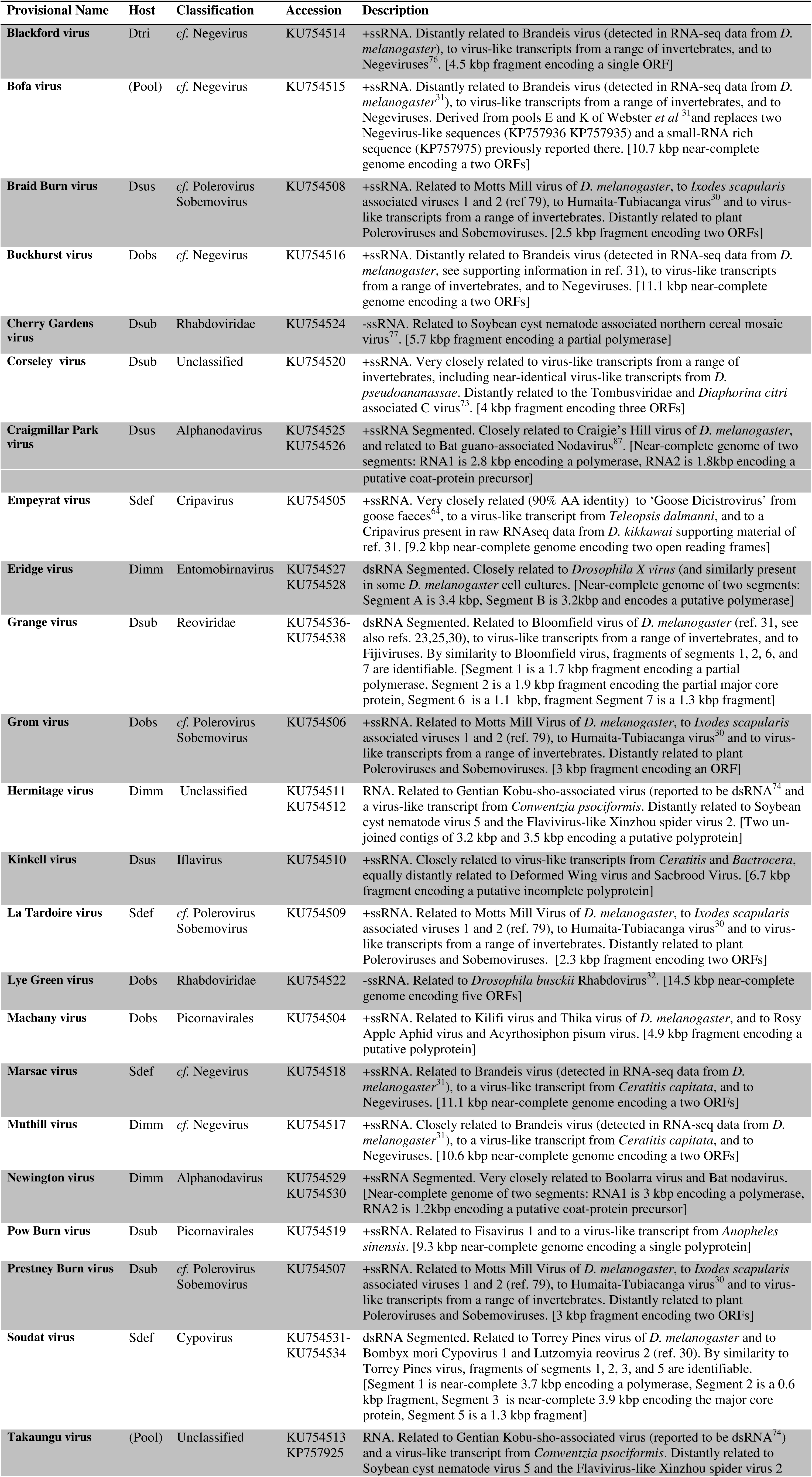
New viruses reported here.

## Supporting Online Data

### Supporting online file 1: Viruses of the Drosophilidae

A comprehensive list of all *Drosophila* viruses reported to date (excluding retroelements) is provided as an xlsx file. Recorded data include: the virus name, its Baltimore classification, the drosophilid hosts in which it has been detected (excluding experimental infections), its year of discovery, its approximate classification, reference Genbank accession identifiers, and citation for its discovery.

### Supporting online file 2: Raw metagenomic assemblies

Compressed fasta files containing the transcriptome assemblies generated for this study (note that as mixed-species (metagenomic) assemblies these cannot be submitted to the NCBI Transcriptome Shotgun Assembly database).

### Supporting online file 3: Alignments used for phylogenetic inference

Compressed fasta-format protein alignments used to infer phylogenetic trees

### Supporting online file 4: Phylogenetic trees

Mid-point rooted maximum likelihood trees, with percentage support marked on nodes for which either tree inference method identified less than 100% support (recorded as ‘SH|bootstrap’), and NCBI accession identifiers for the sequences used to infer the phylogeny.

### Supporting online file 5: Small RNAs (19-30nt) that map to Takaungu virus and Bofa virus from *D. melanogaster*

Bar charts (left) show the size distribution of small RNAs mapping to the positive-sense (above x-axis) and negative-sense (below x-axis) viral strands, and their base composition at the 5' position (red=U, yellow=G, blue=C, green=A). Bar charts (right) show the distribution of 19-30nt reads along the length of the virus contig (blue bars represent reads mapping to the positive strand, red bars represent reads mapping to the negative strand).

## References

1. Huszart T, Imler JL. Drosophila viruses and the study of antiviral host-defense. Advances in Virus Research, Vol 72. Vol 722008:227–265.

2. Xu J, Cherry S. Viruses and antiviral immunity in Drosophila. Dev. Comp. Immunol. Jan 2014;42(1):67–84.

3. Bronkhorst AW, van Rij RP. The long and short of antiviral defense: small RNA-based immunity in insects. Current Opinion in Virology. 2014;7(0):19–28.

4. Chtarbanova S, Imler JL. Innate antiviral immunity in drosophila. Virologie. Sep-Oct 2011;15(5):296–306.

5. Dostert C, Jouanguy E, Irving P, et al. The Jak-STAT signaling pathway is required but not sufficient for the antiviral response of *Drosophila*. Nature Immunology. Sep 2005;6(9):946–953.

6. van Rij RP, Saleh M-C, Berry B, et al. The RNA silencing endonuclease Argonaute 2 mediates specific antiviral immunity in *Drosophila melanogaster*. Genes Dev. November 1, 2006 2006;20(21):2985–2995.

7. Teixeira L, Ferreira A, Ashburner M. The Bacterial Symbiont Wolbachia Induces Resistance to RNA Viral Infections in *Drosophila melanogaster*. PLoS Biology. Dec 2008;6(12):2753–2763.

8. Costa A, Jan E, Sarnow P, Schneider D. The Imd Pathway Is Involved in Antiviral Immune Responses in Drosophila. PLoS ONE. Oct 15 2009;4(10).

9. Nakamoto M, Moy Ryan H, Xu J, et al. Virus Recognition by Toll-7 Activates Antiviral Autophagy in Drosophila. Immunity. 2012;36(4):658–667.

10. Kemp C, Mueller S, Goto A, et al. Broad RNA Interference–Mediated Antiviral Immunity and Virus-Specific Inducible Responses in Drosophila. The Journal of Immunology. January 15, 2013 2013;190(2):650–658.

11. Powell JR. Progress and prospects in Evolutionary Biology: The Drosophila Model. New York: Oxford University Press; 1997.

12. Longdon B, Hadfield JD, Day JP, et al. The Causes and Consequences of Changes in Virulence following Pathogen Host Shifts. PLoS Pathog. 2015;11(3):e1004728.

13. Longdon B, Hadfield JD, Webster CL, Obbard DJ, Jiggins FM. Host Phylogeny Determines Viral Persistence and Replication in Novel Hosts. PLoS Pathog. 2011;7(9):e1002260.

14. Brun G, Plus N. The viruses of Drosophila. In: Ashburner M, Wright TRF, eds. The genetics and biology of Drosophila. New York.: Academic Press; 1980:625–702.

15. Berkalof A, Breglian JC, Ohanessi A. Mise en evidence de virions dans de drosophiles infetees par le virus hereditare Sigma. Comptes Rendus Hebdomadaires Des Seances De L Academie Des Sciences. 1965;260(22):5956-&.

16. L'Héritier PH, Teissier G. Une anomalie physiologique héréditaire chez la Drosophile. C. R. Acad. Sci., Paris. 1937;205:1099.

17. L'Heritier PH. Sensitivity to CO2 in *Drosophila* - A review. Heredity. 1948;2(3):325–348.

18. Plus N, Duthoit JL. Un nouveau virus de *Drosophila melanogaster*, le virus P. Comptes Rendus Hebdomadaires Des Seances De L Academie Des Sciences Serie D. 1969;268(18):2313.

19. Jousset FX, Plus N, Croizier G, Thomas M. Existence chez *Drosophila* de deux groupes de Picornavirus de propriétés sérologiques et biologiques différentes. Comptes Rendus Hebdomadaires Des Seances De L Academie Des Sciences Serie D. 1972;275(25):3043.

20. Plus N, Croizier G, Jousset FX, David J. Picornaviruses Of Laboratory And Wild Drosophila-Melanogaster - Geographical Distribution And Serotypic Composition. Annales De Microbiologie. 1975;A126(1):107-&.

21. Plus N, Croizier G, Duthoit JL, David J, Anxolabehere D, Periquet G. Découverte, chex la Drosophile, de virus appartenant à trois nouveaux groupes. Comptes Rendus Hebdomadaires Des Seances De L Academie Des Sciences Serie D. 1975;280(12):1501-&.

22. Teninges D, Ohanessian A, Richardmolard C, Contamine D. Isolation and biological properties of Drosophila X Virus J. Gen. Virol. 1979;42(FEB):241–254.

23. Teninges D, Ohanessian A, Richard-Molard C, Contamine D. Contamination and Persistent Infection of *Drosophila* Cell Lines by Reovirus Type Particles. In Vitro. 1979;15(6):425–428.

24. Haars R, Zentgraf H, Gateff E, Bautz FA. Evidence for endogenous reovirus-like particles in a tissue culture cell line from Drosophila melanogaster. Virology. 1980/02/01 1980;101(1):124–130.

25. Pasyukova EG, Mukha DV. Reovirus-like Double Stranded RNA Fraction in a *Drosophila melanogaster* Line Containing Individual Second Chromosome from Natural Population. In: Connell CJ, Ralsto DP, eds. Insect Viruses: Detection, Characterization and Roles: Nova Science Publishers, Inc.; 2009:157–164.

26. Jousset FX. Virus extracted from *Drosophila immigrans* inducing a CO_2_ sensitivity syndrom in males of *Drosophila melanogaster*. Comptes Rendus Hebdomadaires Des Seances De L Academie Des Sciences Serie D. 1970;271(13):1141-&.

27. Louis C, Lopez-Ferber M, Comendador M, Plus N, Kuhl G, Baker S. Drosophila S virus, a hereditary reolike virus, probable agent of the morphological S character in Drosophila simulans. Journal of Virology. 1988;62(4):1266–1270.

28. Habayeb MS, Ekengren SK, Hultmark D. Nora virus, a persistent virus in Drosophila, defines a new picorna-like virus family. J. Gen. Virol. Oct 2006;87:3045–3051.

29. Wu Q, Luo Y, Lu R, et al. Virus discovery by deep sequencing and assembly of virus-derived small silencing RNAs. Proceedings of the National Academy of Sciences. January 26, 2010 2010;107(4):1606–1611.

30. Aguiar Eric Roberto Guimarães R, Olmo RP, Paro S, et al. Sequence-independent characterization of viruses based on the pattern of viral small RNAs produced by the host. Nucleic Acids Research. June 3, 2015 2015.

31. Webster CL, Waldron FM, Robertson S, et al. The Discovery, Distribution, and Evolution of Viruses Associated with *Drosophila melanogaster*. PLoS Biol. 2015;13(7):e1002210.

32. Longdon B, Murray GGR, Palmer WJ, et al. The evolution, diversity, and host associations of rhabdoviruses. Virus Evolution. 2015-01-01 00:00:00 2015;1(1).

33. van Mierlo JT, Overheul GJ, Obadia B, et al. Novel *Drosophila* viruses encode host-specific suppressors of RNAi. PLoS Pathog. 2014;10(7):e1004256.

34. Longdon B, Obbard DJ, Jiggins FM. Sigma viruses from three species of Drosophila form a major new clade in the rhabdovirus phylogeny. Proc. R. Soc. B-Biol. Sci. Jan 2010;277(1678):35–44.

35. Longdon B, Wilfert L, Osei-Poku J, Cagney H, Obbard DJ, Jiggins FM. Host-switching by a vertically transmitted rhabdovirus in Drosophila. Biol. Lett. March 30, 2011 2011;7(5):747–750

36. Unckless RL. A DNA Virus of *Drosophila*. PLoS ONE. 2011;6(10):e26564.

37. Piégu B, Guizard S, Yeping T, et al. Genome sequence of a crustacean iridovirus, IIV31, isolated from the pill bug, Armadillidium vulgare. J. Gen. Virol. 2014;95(7): 1585–1590.

38. Joshi NA, Fass JN. Sickle: A sliding-window, adaptive, quality-based trimming tool for FastQ files https://github.com/najoshi/sickle2011.

39. Martin M. Cutadapt removes adapter sequences from high-throughput sequencing reads. EMBnet.journal. May 2011 2011;17(1):10–12.

40. Grabherr MG, Haas BJ, Yassour M, et al. Full-length transcriptome assembly from RNA-Seq data without a reference genome. Nat Biotech. 2011;29(7):644–652.

41. Benson DA, Cavanaugh M, Clark K, et al. GenBank. Nucleic Acids Research. January 1, 2013 2013;41(D1):D36–D42.

42. Camacho C, Coulouris G, Avagyan V, et al. BLAST+: architecture and applications. Bmc Bioinformatics. 2009;10(1):421.

43. Langmead B, Salzberg SL. Fast gapped-read alignment with Bowtie 2. Nat Meth. 2012;9(4):357–359.

44. Brister JR, Ako-adjei D, Bao Y, Blinkova O. NCBI Viral Genomes Resource. Nucleic Acids Research. January 28, 2015 2015;43(D1):D571–D577.

45. Notredame C, Higgins DG, Heringa J. T-coffee: a novel method for fast and accurate multiple sequence alignment1. Journal of Molecular Biology. 2000;302(1):205–217.

46. Larkin MA, Blackshields G, Brown NP, et al. Clustal W and Clustal X version 2.0. Bioinformatics. November 1, 2007 2007;23(21):2947–2948.

47. Lee C, Grasso C, Sharlow MF. Multiple sequence alignment using partial order graphs. Bioinformatics. March 1, 2002 2002;18(3):452–464.

48. Edgar RC. MUSCLE: multiple sequence alignment with high accuracy and high throughput. Nucleic Acids Research. March 1, 2004 2004;32(5):1792–1797.

49. Katoh K, Toh H. Recent developments in the MAFFT multiple sequence alignment program. Briefings in Bioinformatics. July 1, 2008 2008;9(4):286–298.

50. Morgenstern B. DIALIGN 2: improvement of the segment-to-segment approach to multiple sequence alignment. Bioinformatics. March 1, 1999 1999;15(3):211–218.

51. Pei J, Sadreyev R, Grishin NV. PCMA: fast and accurate multiple sequence alignment based on profile consistency. Bioinformatics. February 12, 2003 2003;19(3):427–428.

52. Do CB, Mahabhashyam MSP, Brudno M, Batzoglou S. ProbCons: Probabilistic consistency-based multiple sequence alignment. Genome Research. February 1, 2005 2005;15(2):330–340.

53. Guindon S, Gascuel O. A Simple, Fast, and Accurate Algorithm to Estimate Large Phylogenies by Maximum Likelihood. Systematic Biology. October 1, 2003 2003;52(5):696–704.

54. Le SQ, Gascuel O. An Improved General Amino Acid Replacement Matrix. Molecular Biology and Evolution. July 1, 2008 2008;25(7):1307–1320.

55. Anisimova M, Gil M, Dufayard J-F, Dessimoz C, Gascuel O. Survey of Branch Support Methods Demonstrates Accuracy, Power, and Robustness of Fast Likelihood-based Approximation Schemes. Systematic Biology. October 1, 2011 2011;60(5):685–699.

56. Chung HK, Kordyban S, Cameron L, Dobos P. Sequence analysis of the bicistronic *Drosophila X virus* genome segment A and its encoded polypeptides. Virology. Nov 15 1996;225(2):359–368.

57. Plus N. Endogenous Viruses of Drosophila melanogaster Cell Lines: Their Frequency, Identification and Origin. In Vitro. 1978;14(12): 1015–1021.

58. Duff MO, Olson S, Wei X, et al. Genome-wide identification of zero nucleotide recursive splicing in Drosophila. Nature. 2015;521(7552):376–379.

59. Rodriguez J, Menet Jerome S, Rosbash M. Nascent-Seq Indicates Widespread Cotranscriptional RNA Editing in Drosophila. Molecular Cell. 2012;47(1):27–37.

60. Dasgupta R, Sgro J-Y. Nucleotide sequences of three Nodavirus RNA2's: the messengers for their coat protein precursors. Nucleic Acids Research. September 25, 1989 1989;17(18):7525–7526.

61. Dacheux L, Cervantes-Gonzalez M, Guigon G, et al. A Preliminary Study of Viral Metagenomics of French Bat Species in Contact with Humans: Identification of New Mammalian Viruses. PLoS ONE. 2014;9(1):e87194.

62. Sim SB, Calla B, Hall B, DeRego T, Geib SM. Reconstructing a comprehensive transcriptome assembly of a white-pupal translocated strain of the pest fruit fly Bactrocera cucurbitae. GigaScience. 2015;4(1):1–5.

63. Calla B, Hall B, Hou S, Geib SM. A genomic perspective to assessing quality of mass-reared SIT flies used in Mediterranean fruit fly (Ceratitis capitata) eradication in California. BMC Genomics. 2014;15(1): 1–19.

64. Greninger AL, Jerome KR. Draft genome of goose dicistrovirus. Genome Announcements. 2016;genomeA00068-16(In Press).

65. Reinhardt JA, Brand CL, Paczolt KA, Johns PM, Baker RH, Wilkinson GS. Meiotic Drive Impacts Expression and Evolution of X-Linked Genes in Stalk-Eyed Flies. PLoS Genet. 2014;10(5):e1004362.

66. Geib S, Calla B, Hall B, Hou S, Manoukis N. Characterizing the developmental transcriptome of the oriental fruit fly, Bactrocera dorsalis (Diptera: Tephritidae) through comparative genomic analysis with Drosophila melanogaster utilizing modENCODE datasets. BMC Genomics. 2014;15.

67. Tan Q-Q, Zhu L, Li Y, et al. A *De* NovoTranscriptome and Valid Reference Genes for Quantitative Real-Time PCR in *Colaphellus bowringi*. PLoS ONE. 2015;10(2):e0118693.

68. Stafford-Banks CA, Rotenberg D, Johnson BR, Whitfield AE, Ullman DE. Analysis of the Salivary Gland Transcriptome of *Frankliniella occidentalis*. PLoS ONE. 2014;9(4):e94447.

69. Clarke TH, Garb JE, Hayashi CY, et al. Multi-tissue transcriptomics of the black widow spider reveals expansions, co-options, and functional processes of the silk gland gene toolkit. BMC Genomics. 2014;15(1):1–17.

70. Signor S, Seher T, Kopp A. Genomic resources for multiple species in the Drosophila ananassae species group. Fly. 2013/01/01 2013;7(1):47–56.

71. Hull JJ, Chaney K, Geib SM, et al. Transcriptome-Based Identification of ABC Transporters in the Western Tarnished Plant Bug *Lygus hesperus*. PLoS ONE. 2014;9(11):e113046.

72. Scully ED, Hoover K, Carlson JE, Tien M, Geib SM. Midgut transcriptome profiling of Anoplophora glabripennis, a lignocellulose degrading cerambycid beetle. BMC Genomics. 2013;14(1): 1–26.

73. Nouri S, Salem N, Nigg JC, Falk BW. A diverse array of new viral sequences identified in worldwide populations of the Asian citrus psyllid (Diaphorina citri) using viral metagenomics. Journal of Virology. December 16, 2015 2015.

74. Kobayashi K, Atsumi G, Iwadate Y, et al. Gentian Kobu-sho-associated virus: a tentative, novel double-stranded RNA virus that is relevant to gentian Kobu-sho syndrome. Journal of General Plant Pathology. 2012;79(1):56–63.

75. Shi M, Lin X-D, Vasilakis N, et al. Divergent Viruses Discovered in Arthropods and Vertebrates Revise the Evolutionary History of the Flaviviridae and Related Viruses. Journal of Virology. January 15, 2016 2016;90(2):659–669.

76. Vasilakis N, Forrester NL, Palacios G, et al. Negevirus: a Proposed New Taxon of Insect-Specific Viruses with Wide Geographic Distribution. Journal of Virology. March 1, 2013 2013;87(5):2475–2488.

77. Bekal S, Domier LL, Niblack TL, Lambert KN. Discovery and initial analysis of novel viral genomes in the soybean cyst nematode. J. Gen. Virol. 2011;92(8):1870–1879.

78. Katzourakis A, Gifford RJ. Endogenous Viral Elements in Animal Genomes. PLoS Genet. 2010;6(11):e1001191.

79. Tokarz R, Williams SH, Sameroff S, Leon MS, Jain K, Lipkin WI. Virome Analysis of *Amblyomma americanum*, *Dermacentor variabilis*, and *Ixodes scapularis* Ticks Reveals Novel Highly Divergent Vertebrate and Invertebrate Viruses. Journal of Virology. Oct 2014;88(19):11480–11492.

80. Li C-X, Shi M, Tian J-H, et al. Unprecedented genomic diversity of RNA viruses in arthropods reveals the ancestry of negative-sense RNA viruses. eLife. 2015-01-29 13:52:37 2015.

81. Ballinger MJ, Bruenn JA, Hay J, Czechowski D, Taylor DJ. Discovery and Evolution of Bunyavirids in Arctic Phantom Midges and Ancient Bunyavirid-Like Sequences in Insect Genomes. Journal of Virology. August 15, 2014 2014;88(16):8783–8794.

82. Han YH, Luo YJ, Wu QF, et al. RNA-Based Immunity Terminates Viral Infection in Adult Drosophila in the Absence of Viral Suppression of RNA Interference: Characterization of Viral Small Interfering RNA Populations in Wild-Type and Mutant Flies. Journal of Virology. Dec 2011;85(24): 13153–13163.

83. Flynt A, Liu N, Martin R, Lai EC. Dicing of viral replication intermediates during silencing of latent Drosophila viruses. Proceedings of the National Academy of Sciences of the United States of America. Mar 2009;106(13):5270–5275.

84. Zambon RA, Vakharia VN, Wu LP. RNAi is an antiviral immune response against a dsRNA virus in *Drosophila melanogaster*. Cell Microbiol. May 2006;8(5):880–889.

85. van Cleef KWR, van Mierlo JT, Miesen P, et al. Mosquito and Drosophila entomobirnaviruses suppress dsRNA‐ and siRNA-induced RNAi. Nucleic Acids Research. July 29, 2014 2014;42(13):8732–8744.

86. Bronkhorst AW, van Cleef KWR, Venselaar H, van Rij RP. A dsRNA-binding protein of a complex invertebrate DNA virus suppresses the Drosophila RNAi response. Nucleic Acids Research. October 29, 2014 2014;42(19):12237–12248.

87. Li L, Victoria JG, Wang C, et al. Bat Guano Virome: Predominance of Dietary Viruses from Insects and Plants plus Novel Mammalian Viruses. Journal of Virology. Jul 2010;84(14):6955–6965.

